# Prevalence of dual clarithromycin and metronidazole resistance in *Helicobacter pylori* estimated by molecular methods in Santiago, Chile

**DOI:** 10.1101/236976

**Authors:** Patricio Gonzalez-Hormazabal, Maher Musleh, Susana Escandar, Hector Valladares, Enrique Lanzarini, V. Gonzalo Castro, Lilian Jara, Zoltan Berger

## Abstract

**Background:** Current available treatments for *Helicobacter pylori* eradication are chosen according to local clarithromycin and metronidazole resistance prevalence. The aim of this study was to estimate, by means of molecular methods, both clarithromycin and metronidazole resistance in gastric mucosa from patients infected with *H.pylori*.

**Methods:** A total of 191 DNA samples were analyzed. DNA was purified from gastric mucosa obtained from patients who underwent an upper gastrointestinal endoscopy at an university hospital from Santiago, Chile, between 2011 and 2014. *H.pylori* was detected by real-time PCR. A 5’exonuclease assay was developed to detect A2142G and A2143G mutations among *Hpylori*-positive samples. *rdxA* gene was sequenced in samples harboring A2142G and A2143G mutations in order to detect mutations that potentially confer dual clarithromycin and metronidazole resistance.

**Results:** Ninety-three (93) out of 191 DNA samples obtained from gastric mucosa were H. pylori-positive (48.7%). Clarithromycin-resistance was detected in 29 samples (31.2% [95%CI 22.0%-41.6%]). The sequencing of *rdxA* gene revealed that two samples harbored truncating mutations in *rdxA*, one sample had an in-frame deletion, and 11 had amino acid changes that likely cause metronidazole resistance.

**Conclusions:** We estimated a prevalence of clarithomycin-resistance of 31.8% in Santiago, Chile. The proportion of dual clarithromycin and metronidazole resistance could be, at least, 15.0%. Our results require further confirmation. Nevertheless, they are significant as an initial approximation in re-evaluating the guidelines for *H.pylori* eradication currently used in Chile.

## 1. Background

*Helicobacter pylori* is a gram-negative bacillus that colonizes the gastric mucosa. The infection is associated with both gastric and extragastric diseases such as peptic ulcer disease, gastric atrophy, gastric adenocarcinoma, gastric MALT (mucosa-associated lymphoid tissue) lymphoma, and iron deficiency anemia. The evidence strongly suggests that *H.pylori* eradication is beneficial in treatment or prevention of those diseases [1]. One treatment for *H.pylori* eradication is the triple-therapy that comprises clarithromycin, amoxicillin, and proton pump inhibitor (PPI), nevertheless, triple-therapy is not recommended in regions with high clarithomycin resistance rate [1]. Global claritrhomycin-resistance rates in adults are contrasted among various countries, from 1% in The Netherlands to 37.6% in Italy [2]. In light of this observation, the last Maastricht V/Florence consensus on management of *H.pylori* [1] recommends that in high-clarithromycin resistance regions (>15% resistance prevalence), quadruple therapy is necessary. If dual clarithromycin and metronidazole resistance is >15%, bismuth-containing quadruple therapy is the recommended first-line treatment.

Clarithromycin belongs to the class of macrolide antibiotics. The mechanism of action is the inhibition of protein synthesis of bacteria by binding to the 23S ribosomal subunit (23S rRNA). Mutations in the peptidyltransferase region encoded in domain V of 23S rRNA are responsible for macrolide resistance [3]. Three mutations in the 23S rRNA gene account for nearly all clarithromycin-resistant (ClaR) strains: A2143G, A2142G and A2142C to a small extent [3]. Standard method (antibiogram), after culture or molecular tests, can be used to detect *H.pylori* and clarithromycin resistance directly from the gastric biopsy [1]. Epidemiological studies evaluating regional clarithromycin resistance have been conducted based on detection of A2142G and A2143G in DNA isolated from gastric biopsies [4,5].

Commercial tests detecting both mutations are available [3], performing at a sensitivity rate of >94% and a specificity rate of >98.5% compared to the culture susceptibility test. Recently, Tamayo et al. [6] found a high concordance between Etest for clarithromycin and 23S rRNA mutations. To date, only Garrido and Toledo [7] have evaluated primary resistance of *H.pylori* to clarithromycin in clinical isolates from Santiago, Chile. They found primary resistance in 10 out of 50 isolates by agar dilution. Resistant isolates harbor A2142G or A2143G mutations.

Metronidazole is a nitroimidazole synthesized as an inactive antibiotic prodrug. The bactericidal effect of the drug is mainly explained by the cytotoxic effect of molecules resulting from a reduction of metronidazole intracellularly in facultative anaerobic bacteria. Reduction of metronidazole in *H.pylori* is mediated mainly by a nonessential oxygen-insensitive NADPH nitroreductase encoded by the *rdxA* gene. NADPH-flavin-oxidoreductase (FrxA) is also involved in the reduction of metronidazole [3]. Various authors found that metronidazole resistance arises from inactivating mutations in *rdxA* found only in resistant isolates, and missense mutations have also been involved in metronidazole resistance [8]. On the other hand, the role of inactivating mutations in the *frxA* gene is controversial since truncating mutations have been found in sensitive isolates, and some studies report that *frxA* mutations alone may not be enough to confer resistance [8]. Vallejos et al. [9] studied resistance to clarithromycin and metronidazole in 50 isolates obtained from patients in Santiago, Chile. Forty-five and twenty percent respectively were resistant, resulting in 13.2% of the prevalence of dual clarithromycin and metronidazole resistance.

The aim of the present study was to estimate clarithomycin resistance in 93 clinical isolates from Santiago, Chile. To do so, we developed a test to detect A2142G and A2143G mutations in 23S gene by 5’nuclease assay. We also aimed to detect mutations in the *rdxA* gene among ClaR isolates.

## 2. Methods

### 2.1. Subjects

Subjects invited to participate were patients who underwent a physician-requested upper gastrointestinal endoscopy, at the Department of Gastroenterology at the University of Chile Clinical Hospital between July 2011 and July 2014. Participants were asked to donate a sample of gastric mucosa obtained from antrum and corpus. This study was approved by the Ethical Committee of the following institutions: University of Chile School of Medicine (#023/2011), and University of Chile Clinical Hospital (#029/2011). The study was performed in accordance with the Declaration of Helsinki. DNA was isolated from gastric mucosa using FavorPrep Tissue DNA Extraction Mini Kit (Favorgen Biotech Corp, Taiwan, China). Samples were identified as *H.pylori*-positive or *H.pylori*-negative by detection of the 16S rRNA gene by real-time PCR as described by Kobayashi et al. [10].

### 2.2. Analysis of 23S rRNA mutations

A segment of the 23S rRNA gene from position 1911 to 2200 was amplified by PCR using primers described by Agudo et al. [11], and sequenced by Sanger sequencing (Service provided by Macrogen Inc., Korea). A 5’exonuclease assay was developed to detect A2142G and A2143G mutations. We designed primers and probes for each allele in highly conserved regions from position 1911 to 2200, comparing sequences obtained from 44 randomly selected *Hpylori*-positive subjects recruited for this study. Both mutations were tested in separate assays. Probes detecting 2142G and 2143G alleles were dual-labeled with 5’FAM-3’BHQ1, and the probe complementary to the 2142A - 2143A allele was dual-labeled with 5’HEX-3’BHQ1. Primers and probes were synthesized at Macrogen Inc. (Korea). The 5’exonuclease was carried out in a StepOne Real Time PCR system (Applied Biosystems, USA) using 5X HOT FIREPol Probe qPCR Mix Plus (ROX) (Solis BioDyne, Estonia) according to manufacturer’s directions. Primer and probe sequences are available by request.

### 2.3. Detection of mutations in rdxA gene

The sequence of *rdxA* was obtained from samples harboring 23S 2142G or 2143G. Primers for PCR amplification of *rdxA* (F: CRTTAGGGATTTTATTGTATGC R: CTCTTRCCCAAWGCGATC) were designed with AliView 1.18 for Linux in conserved regions after comparing 83 sequences of *H.pylori* obtained from GenBank. PCR was performed using Q5 Hot Start High Fidelity DNA Polymerase (New England Biolabs, USA) according to manufacturer’s directions, using an annealing temperature of 56°C. Electrophoretograms were obtained by Sanger sequencing (Service provided by Macrogen Inc., Korea).

## 3. Results

A total of 191 subjects were included in this study, 124 were female (64.9%) and 67 male (35.1%). The median age was 46 years (range 19 years to 82 years). None of them was treated for *H.pylori* eradication. Ninety-three (93) out of 191 DNA samples obtained from gastric mucosa were *H.pylori*-positive (48.7%). Clarithomycin resistance was detected using a 5’exonuclease assay in 29 samples (31.2% [95%CI 22.0%-41.6%]). Of them, 9 harbored 2142G (31.0%) and 20 (69.0%) harbored 2143G. All ClaR samples were confirmed by Sanger sequencing. Both Sanger sequencing and 5’exonuclease allowed us to detect heteroresistance, i.e., the coexistence of resistant and susceptible bacterial strains in the same patient [12]. Sixteen (16) out of 93 *H.pylori* subjects were homoresistant to clarithromycin (17.2%). In addition, we detected heteroresistance in 13 out of 93 patients (14.0%).

The *rdxA* gene was sequenced in 28 ClaR samples. One ClaR sample was not available at the time of analysis. A nonsense substitution (H53stop) was found in one sample; and an insertion of A at nucleotide 186, leading to a frameshift resulting in N73stop, was found in a different sample. Therefore, two samples harbored truncating mutations in *rdxA*. One sample had an in-frame deletion of 39 nucleotides which causes the deletion of amino acids 80 to 92 (ASALMVVCSLKPS). Table 1 shows the amino acid variations in *rdxA* found in ClaR samples.

## 4. Discussion

A total of 29 out of the 93 *H.pylori* positive samples resulted resistant to clarithomycin (31.2%). This estimation is higher than the 20% of ClaR reported by Garrido and Toledo in Chile [7]. Taken together, ClaR resistance reported for Chile is above the suggested 15% threshold to abandon triple-therapy, according to the recommendation of Maastricht V/Florence consensus [1].

Among 28 ClaR samples, two of them harbored truncating mutations and one an inframe deletion in *rdxA*, and therefore are likely resistant to metronidazole. A total of 22 amino acid substitutions was found. Thirteen of them (Q6H, T31E, H53R, L62V, R90K, H97T, H97Y, G98S, A118T, V123T, V172I, E175Q, A183V) have been previously observed in metronidazole-resistant (MtzR) as well as in metronidazole-sensitive strains [13–17]. Therefore, those substitutions probably do not confer resistance to metronidazole. Four substitutions (R16H, A67V, S88P and G162R) were found in 8 samples, and likely cause resistance. R16H and S88P have been observed only in MtzR strains [13–15]. In fact, Arg 16 is one of the residues of RdxA that interacts with the cofactor flavin mononucleotide (FMN) [18]. A67V corresponds to the replacement of a small alanine by a bulky valine in the protein core, and is associated with resistance [8,19]. Analysis from the crystallographic data of RdxA reveals that G162 is involved in FMN binding [18].

Five amino acid changes in RdxA (T49M, E75Q, V151L, P166A and P166S) have not been described previously in the literature. None of them have been observed as an important amino acid for the RdxA function. We predicted the effect of those substitutions using the crystallographic data of RdxA protein (Protein Data Bank accession number 3QDL) in the I-Mutant Suite server [20], which estimates protein stability changes (expressed as delta delta G values) upon single-point mutations from the protein structure. Delta-delta G values and the prediction were as follows: T49M = -0.41 (neutral stability), E75Q = -0.78 (large decrease of stability), V151L = -0.44 (neutral stability), P166A = -1.28 (large decrease in stability), P166S = -1.50 (large decrease in stability). Thus E75Q (1 sample), P166A (1 sample), and P166S (1 sample) could confer resistance to metronidazole. This conclusion deserves to be confirmed by additional studies. Taken together, 14 out of 93 analyzed samples (15.1% [95%CI 7.8% - 22.3%]) contain *H.pylori* strains likely resistant for both metronidazole and clarithromycin.

## 5. Conclusions

We estimated a prevalence of ClaR of 31.2% in Santiago, Chile. The proportion of dual clarithromycin and metronidazole resistance could be, at least, 15.0%. Our results require further confirmation. Nevertheless, they are significant as an initial approximation in reevaluating the triple-therapy for *H.pylori* eradication currently used in Chile.

## 6. List of abbreviations

PPI: Proton pump inhibitor
ClaR: Clarithromycin-resistant

## 7. Declarations

### 7.1. Ethics approval and consent to participate

This study was approved by the Ethical Committee of the following institutions: University of Chile School of Medicine (#023/2011), and University of Chile Clinical Hospital (#029/2011). The study was performed in accordance with the Declaration of Helsinki.

### 7.2. Availability of data and materials

The data that support the findings of this study are available from the corresponding author upon reasonable request.

### 7.5. Authors’ contributions

PGH conceived, designed the study and drafted the manuscript. PGH, MM and ZB were responsible for analysis and interpretation of data. VGC and LJ performed molecular analysis. MM, SE, HV and EL were involved in acquisition of samples and data. All authors read and approved the final manuscript.

## 7.6. Acknowledgement

We would like to acknowledge Paul Zuckerman for his help in proofreading the manuscript.

